# Uncoupling of dysgranular retrosplenial “head direction” cells from the global head direction network

**DOI:** 10.1101/062042

**Authors:** Pierre-Yves Jacob, Giulio Casali, Laure Spieser, Hector Page, Dorothy Overington, Kate Jeffery

**Author notes:** Correspondence to: Pierre-Yves Jacob or Kate Jeffery.

## Abstract

Spatial cognition is an important model system with which to investigate how sensory signals are transformed into cognitive representations. Head direction cells, found in several cortical and subcortical regions, fire when an animal faces a given direction and express a global directional signal which is anchored by visual landmarks and underlies the “sense of direction”. We investigated the interface between visual and spatial cortical brain regions and report the discovery that a population of neurons in the dysgranular retrosplenial cortex, which we co-recorded with classic head direction cells in a rotationally symmetrical two-compartment environment, were dominated by a local visually defined reference frame and could be decoupled from the main head direction signal. A second population showed rotationally symmetric activity within a single sub-compartment suggestive of an acquired interaction with the head direction cells. These observations reveal an unexpected incoherence within the head direction system, and suggest that dysgranular retrosplenial cortex may mediate between visual landmarks and the multimodal sense of direction. Importantly, it appears that this interface supports a bi-directional exchange of information, which could explain how it is that landmarks can inform the direction sense while at the same time, the direction sense can be used to interpret landmarks.

## Main text

The brain’s construction of knowledge involves a transformation between local, perceptual and more global, cognitive representations. In the spatial domain, directional orientation involves using (mainly visual) landmarks to establish and maintain a “sense of direction”. Central to this process are the head direction (HD) cells (Taube, 2007), which fire when the animal faces a particular direction and are oriented by landmarks (Taube et al., 1990). Retrosplenial cortex (RSC) is a cortical head direction cell region (Chen et al., 1994a, Chen et al., 1994b; Cho and Sharp, 2001): it is directly connected to visual cortex (van Groen and Wyss, 1990, van Groen and Wyss, 1992; Van Groen and Wyss, 2003) and may form the link between visual landmarks and the direction sense. To investigate this link, we recorded RSC HD cells in two connected but visually rotated compartments distinguished only by their odour, to see how the system resolves the conflict between the visual inputs and the global, directional signal.

Rats moved freely between two rectangular compartments through a central doorway (Fig. 1a; Supplementary Data Fig. 1). Each compartment formed a local reference frame that was polarized by visual cues (a cue card, plus the doorway) but was reversed in orientation compared to the other compartment. The overall two-fold rotational symmetry of this arrangement was broken by scenting one compartment with lemon and one with vanilla. Recordings were made from RSC in 5-trial sessions with the door open (trials 1, 2 and 5) or closed (trials 3 and 4). Of the 1090 RSC neurons we recorded (Table 1), 96 neurons (9%) were identified as HD cells (selection criteria in Methods), with classic unipolar directional tuning curves that were oriented by the olfactory cues (Supplementary Data Fig. 2; see Supplementary information) and which maintained consistency across compartments even if the start compartment was randomly chosen (Fig. 1b). Thus, odour context can provide directional information.

**Table 1:** Summary of the cell distributionMethods Subjects

|  | **Rat 599** | **Rat 602** | **Rat 628** | **Rat 636** | **Total (n)** | **Total (%)** |
| --- | --- | --- | --- | --- | --- | --- |
| No. recording sessions | 28 | 25 | 16 | 16 | 85 |  |
| Total cells | 591 | 411 | 48 | 40 | 1090 |  |
| HD cells | 35 | 44 | 4 | 13 | 96 | 9 |
| All bipolar cells | 76 | 16 | 21 | 3 | 116 | 11 |
| Between-compartment | 30 | 5 | 10 | 1 | 46 | 4 |
| Within-compartment | 46 | 11 | 11 | 2 | 70 | 6 |

Importantly, we also recorded many cells having two well-defined head-direction tuning curves separated by 180° (Fig. 1c; see Supplementary Data Fig. 3 for more examples). Henceforth, we call these “bipolar HD cells” to distinguish them from the unipolar HD cells. To quantify our observations we derived a measure of bipolarity, the “flip score” (Supplementary Data Fig. 4), which we used to identify 116 bipolar cells (11% of the total). Of these neurons, 46 (40% of bipolar cells) were compartment-specific (Fig. 1c and Supplementary Data Fig. 3). Thus, cells fired in a specific direction in a given compartment, and reversed their firing by 180 degrees when the rat crossed into the second compartment. After closing the door, these cells typically now exhibited a single unipolar tuning curve in each compartment, recovering the bipolar pattern when the door re-opened.

We used several methods to rule out that the bipolar tuning curves might have been artefact (Supplementary Data Fig. 5). However, the definitive refutation is that we recorded ordinary, unipolar HD cells simultaneously with the bipolar ones (e.g., Fig. 1d; see Supplementary Data Fig. 6 for additional examples). Bipolar cells also followed the cues more reliably than unipolar cells (Supplementary Data Fig. 7), reinforcing the dissociation between the two cell types. This observation reveals a major incoherence within the head direction system and suggests that some dysgranular retrosplenial neurons lie outside the main head direction network.

**Figure 1:**
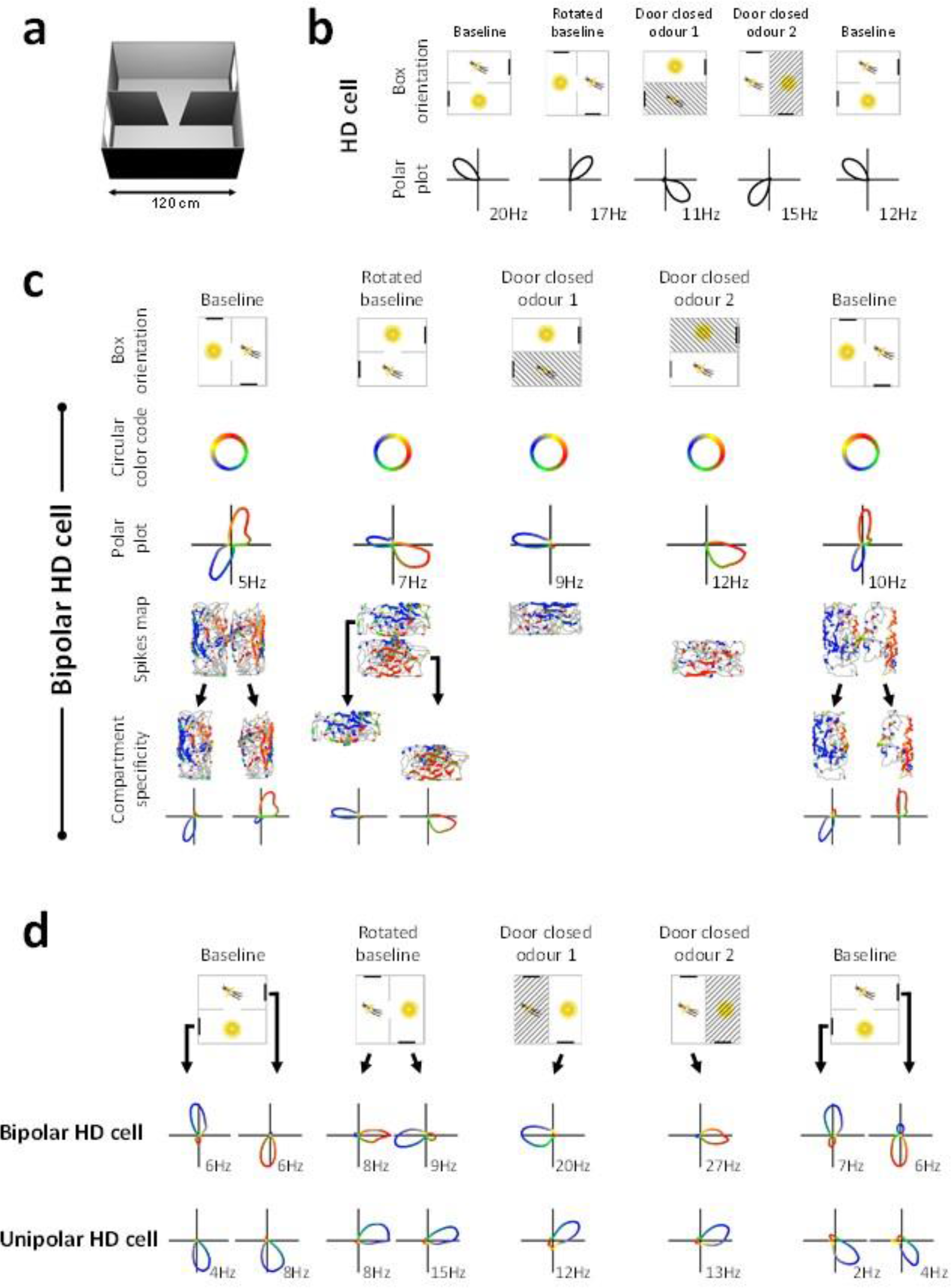
Two types of directional encoding by RSC neurons. **a**, Schematic of the two-compartment apparatus, **b**, Example of an RSC HD cell. Top: schematic diagram of the apparatus orientation (cross-hatching = unavailable compartments). Bottom: polar plotsof the cell’s firing rate as a function of head direction (no. = peak rate), **c**, Example of an RSC bipolar HD cell. Top row: apparatus orientation. Second row: circular colour key. Third row: tuning curves color-coded accordingtothe colourkey. Fourth row: path of the rat (grey lines) with overlying spikes directionally color-coded as per the colour key. Bottom row: tuning curves from each sub-compartment in isolation, **d**, Simultaneous recordings of a bipolar and unipolar HD cell, color-coded as before, revealing the dissociation in tuning curve direction.

We looked for bipolar firing patterns in two other HD-cell-containing structures, the postsubiculum (PoS) and the anterior thalamus (ADN), but found only classic unipolar HD cells (Fig. 2a, Table 1, see Supplementary Data Fig. 4 for bipolar vs unipolar analysis). Within RSC, histology showed a distribution of unipolar HD cells across both granular and dysgranular RSC while bipolar HD cells occurred exclusively only in dysgranular (Fig 2b and Supplementary Data Fig. 8). Thus, these cells occur only in a restricted region of the HD circuit.

The sudden reversal of the directional firing of bipolar HD cells as rats traversed the doorway suggests an influence of visual cues, since these also rotated between compartments. However, 12 bipolar HD cells recorded in the dark maintained their bipolar pattern (average flip score for both ‘darkness’ and ‘rotated darkness’ trials = 1.06 ±0.4; Supplementary Data Fig. 9 a-c). Thus, at least once the environment is familiar, other cues including olfactory and tactile can evidently mediate the influence of environmental cues on firing. We also recorded 9 bipolar HD cells in an open field, and saw a complete loss of the bipolar pattern (average flip score = −0.17 ±0.18; Supplementary Data Fig. 9 d, e), further suggesting that it arises from environment structure rather than intrinsic cell properties.

**Figure 2:**
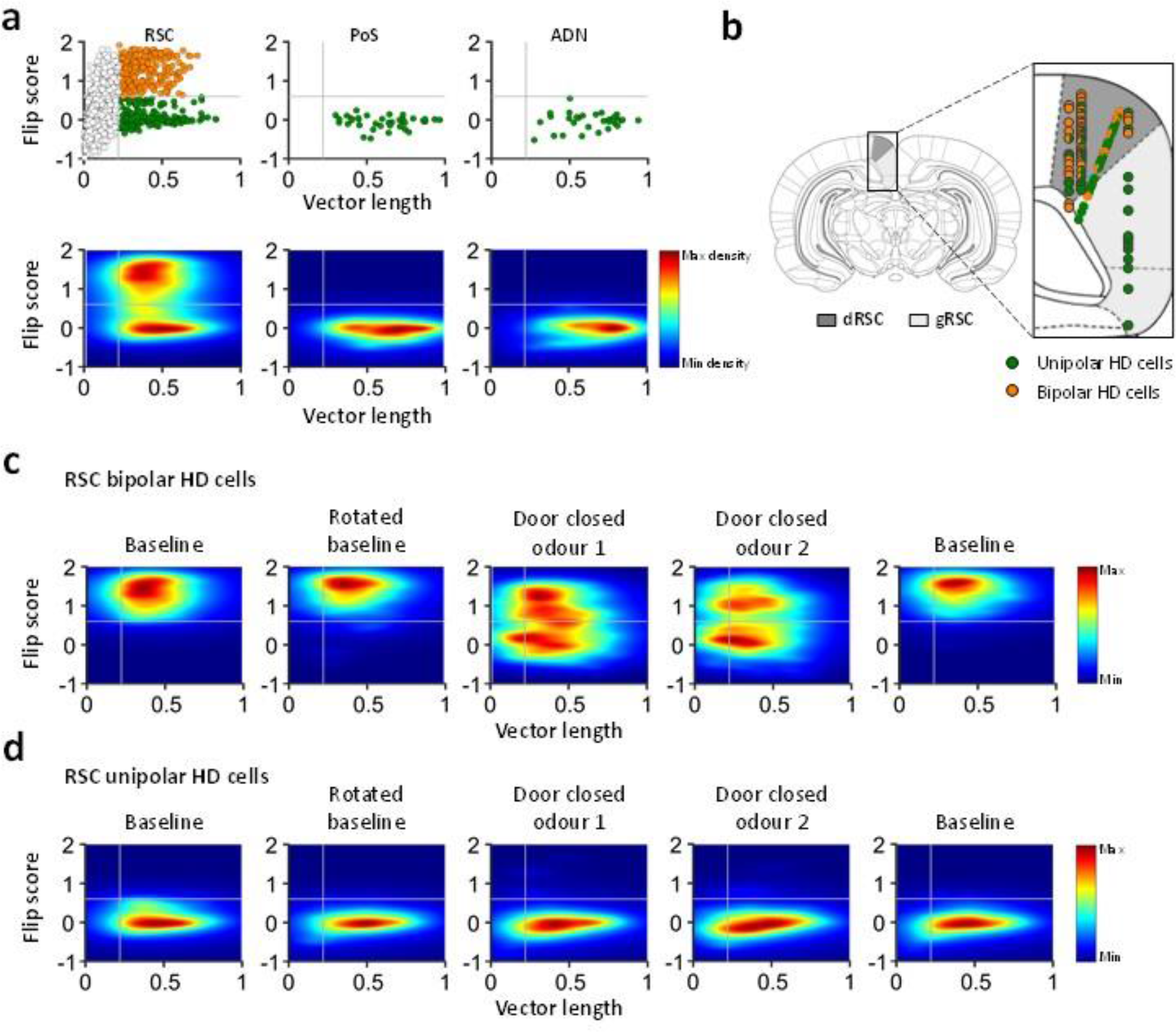
Bipolar cells are specific to dysgranular RSC. **a**, Distribution of flip score as a function of directional tuning (angle doubled Rayleigh vector length) for 1090 RSC cells in trial 1. For PoS and ADN cells, we used the subset of neurons that reach the angle-doubled vector length threshold 0.22. Top, scatter plot shows that RSC contains both bipolar (orange) and unipolar (green) HD cells, while PoS and ADN contain exclusively HD cells. Bottom, density plots of the same data revealing two clear clusters in RSC. **b**, Schematic diagram (adapted from Paxinos and Watson, 2007) of anatomical recording sites of bipolar (orange) and unipolar (green) HD cells within the dysgranular (dRSC, dark grey) and granular (gRSC, light grey) RSC. **c**,**d**, Density plots of RSC bipolar (**c**) and unipolar (**d)** cells across 5 recording trials, collapsed across cells/sessions. In single-compartment trials (3 and 4), some bipolar cells (**c**) acquired a low flip score (resembled HD cells) and some retained a high flip score indicating within-compartmental bipolarity. HD cells (**d**) maintained a low flip score (single tuning curve) across all 5 trials.

As well as the cells described above that reversed firing directions between compartments, we also found that 70/116 bipolar HD cells continued to express a bipolar pattern *within* each sub-compartment (Fig. 2c and Fig. 3a, b). However in individual cells, the dominant (larger) peak reversed direction between compartments (Fig. 3 c) indicating continued influence of the polarising landmarks as well. We investigated whether these neurons possessed a unipolar tuning curve that reversed periodically throughout the trial, or simultaneous opposing tuning curves. In the reversing-unipolar case, the number of spikes overall would be the same as for a true unipolar HD cell, while the spread of firing rates across epochs in the head faced in a given tuning curve direction would be increased, due to the quiescent periods accompanying state reversal. In fact, the spike count was 37% higher than for HD cells, and 42% higher than for between-compartment bipolar cells (Fig 3d; [F(2,211) =4.282, *p* < 0.05], both differences being significant (Tukey HSD corrected). Spread of firing rates was no different for the three cell types (Fig 3e; [F(2,211) =2.711, n.s.]), and inter-spike interval (ISI) analysis showed no increase in long intervals suggestive of state reversals; moreover, analysis of the ISI histogram decay time-to-half-peak showed no difference between cell types [F(2,157) = 1.33, NS]. Taken together, these observations suggest that the cells have true dual tuning curves (one larger, one smaller) rather than single, periodically reversing ones.

**Figure 3:**
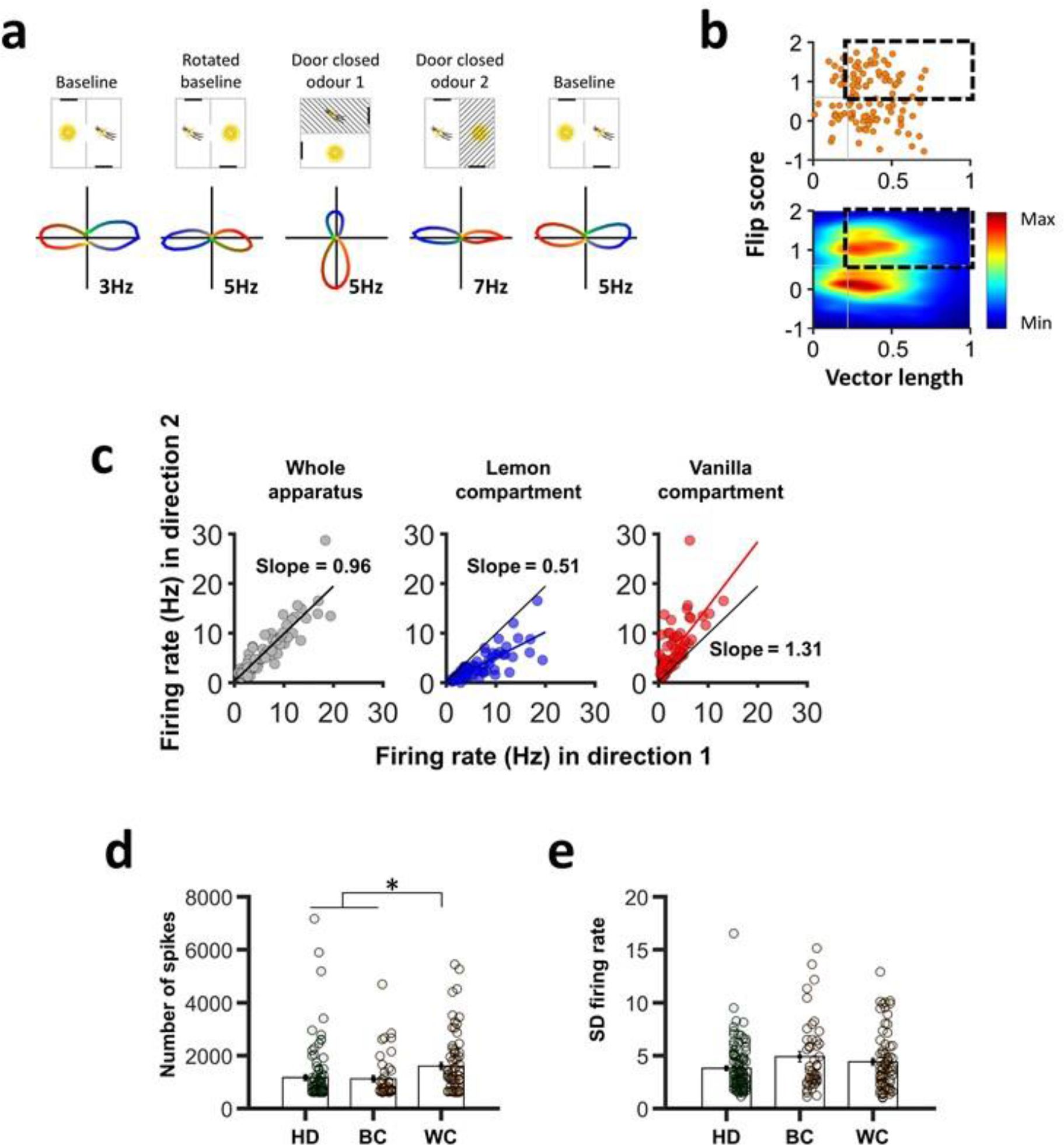
Within-compartmental bipolar activity. **a**, A bipolar cell with bi-directional firing in the single-compartmenttrials. **b**, 60% of bipolarcells showed a within-compartmental bipolarity, **c**, Asymmetry of firing rateforthe two peaks in the within-compartment bipolarcells. Direction 1 was defined as the direction of the biggest peak in the lemon compartment, and direction 2 the opposite direction. The size asymmetry reversed direction between compartments, suggesting a spatial influence. **d**,**e** Analysis of spike count (d) revealed an increase in spikingforthe within-compartment bipolarcells (WC; * = p < 0.05) compared with the between-compartment bipolarcells (BC) and the head direction cells (HD)․. However, firing rate spread (e) was not different between cell types. Solid lines show mean and s.e.m․.

What could cause this within-compartment bipolar pattern? Because bipolar HD cells have not previously been reported in RSC, we considered it likely that these firing patterns arise as a consequence of the rat’s previous exploration of the two-compartment space. We hypothesize that it is due to acquisition of inputs from HD cells (Fig. 4). Because the visual landmarks in our unusual apparatus have two opposing relationships to the HD signal, if an initially landmark-sensitive cell could acquire (by Hebbian learning) additional inputs from co-active HD cells, then it would associate with two sets of HD cells having opposing firing directions, one set from each compartment. This could explain why one tuning curve was slightly stronger (because it was additionally driven by the landmark), and why the direction of the stronger tuning curve reversed between compartments (because the landmarks

Finally, we looked at the temporal and spatial properties of the three different types of head direction cells. ADN neurons had higher peak rates and shorter inter-spike intervals (Supplementary information) but no other salient differences were found. We also looked at movement and spatial correlates, as RSC neurons have been reported to have conjunctive coding properties (Cho and Sharp, 2001; Alexander and Nitz, 2015). There was no correlation of neuronal firing rate with either linear or angular head velocity for either cell type. They may, therefore, be neurons of a similar type that simply vary in the pattern of inputs from landmarks vs true HD cells.

**Figure 4:**
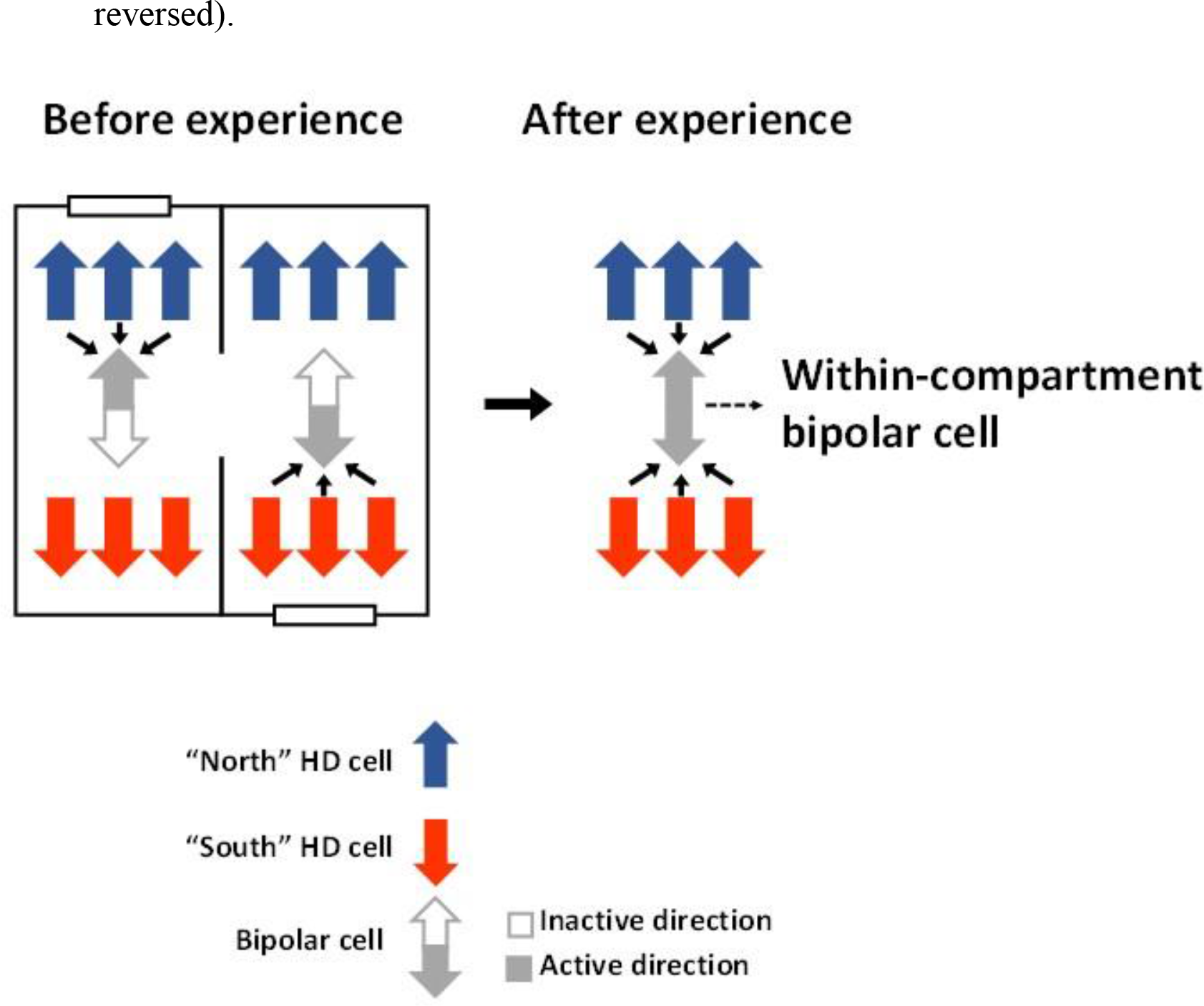
Hypothesized formation of within-compartment bipolar cell. A rat exploring the two compartment box undergoes activation of both head direction cells and bipolar cells. The bipolar cell, because of its reversal, is co active with two opposing sets of HD cells. The consequence is that the neuron now has inputs from both sets, and can be driven to firing threshold in both directions.

What could be the function of bipolar HD cells? Because they are evidently landmark-driven, and yet have the properties of HD cells rather than of visual cells (such as firing in the same direction across the apparatus, and maintaining firing in the dark), it may be to mediate bi-directionally between visual landmarks and the global HD signal. A network of neurons having varying degrees of coupling with landmarks vs. with the global HD network could, in a bootstrapping manner, simultaneously compute both head direction relative to landmarks, and landmarks relative to head direction. Thus, the network could learn which cues are the directionally stable ones that can function as landmarks to define a local spatial reference frame. This hypothesis is consistent with evidence from gene activation studies suggesting a role for dysgranular RSC in spatial tasks requiring light (i.e. vision) but not darkness (Cooper et al., 2001; Pothuizen et al., 2009), human neuroimaging studies showing a role for RSC in monitoring landmark stability (Auger et al., 2012) and local reference frames (Marchette et al., 2014), and from recording studies suggesting plasticity in the connections of landmarks to the HD network (Knight et al., 2014).

Our observation of cells that resemble HD cells in every respect, but are influenced by landmarks independent of the main HD signal, suggests that these are neurons whose function is to mediate between more perceptual and more cognitive parts of the spatial circuit. Understanding the interaction between these cell types at a network level may yield important insights not just about spatial cognition, but about sensory integration and knowledge representation more generally.

## Acknowledgments

The data reported here are held in a UCL data repository and are available via direct request to the authors. This work was supported by grants from the Medical Research Council (G1100669) and Wellcome Trust (103896AIA) to K.J.

## Authors contribution

P.Y.J. and K.J. designed the study, P.Y.J. and D.O. performed surgeries and recordings, P.Y.J., L.S., G.C. and H.P. analysed data. All authors interpreted data and
discussed results. P.Y.J. and K.J. wrote the manuscript. All authors commented and edited the manuscript. Authors declare no conflict of interest.

## Methods

### Subjects

All procedures were carried out under the auspices of a Home Office license according to the Animals (Scientific Procedures) Act 1986. Nine adult male Lister Hooded rats weighing 300-350g were individually housed and mildly food restricted (to 90% free-feeding weight). The animals were weighed and checked daily.

### Microdrives and surgery

Recordings were made using tetrodes, each composed of four twisted 25 p,m or eight twisted 17 μm polyimide-coated platinum-iridium (90%/10%) wires (California Fine Wire, CA), attached to an Axona microdrive (Axona Ltd, Herts, UK). Under isoflurane anaesthesia the electrodes were chronically implanted in the left or right hemisphere. For 4 rats the electrodes were implanted in the RSC aimed at the granular subregion (n = 1 rat; co-ordinates in mm from Bregma: AP: −5.5, ML: ±0.4, DV: 0.4), or dysgranular subregion (n = 3 rats; AP: −5.5, ML: ±1.0, DV: 0.4). For three rats the electrodes were aimed at postsubiculum (AP: −7.5, ML: ±3.2, DV: −2.0) and for two rats they were aimed at the anterodorsal thalamic nucleus (ADN; AP: −1.8, ML: ±1.4, DV: −3.9). After the surgery, rats received meloxicam mixed with condensed milk for three days for post-operative analgesia, and were given at least 7 days to recover before the experiment began.

### Apparatus

The recording apparatus (Fig. 1a), Supplementary Data Fig. 1a) was a 120 × 120 cm square box with 60 cm high walls, isolated from the rest of the room by a cylindrical curtain 260 cm in diameter. The box was divided into two equal rectangular sub-compartments by a 60 cm high wall, in the centre of which was an aperture, 10 cm at the base and 15 cm at the top, which allowed free movement between compartments. The compartments were each polarized by a 20 cm wide × 40 cm high plastic white cue card attached to the short wall and located to the left when facing the doorway. Walls and floor were covered with black vinyl sheets, allowing the experimenter to wipe each compartment with a sponge moistened with lemon or vanilla food flavouring so as to allow the rat to distinguish the compartments. Six circularly arranged ceiling lights lighted the box and a non-tuned radio was fixed to the ceiling in a central position relative to the box, producing a background noise >70 dB to mask un-controlled directional sounds. For recordings made in darkness, the cue cards were removed so as to minimize focal olfactory/tactile landmarks.

### Recording setup and procedure

For recording, the microdrive connector was attached to the headstage/recording cable through which the signals from each wire were amplified, filtered, digitized (48 kHz) and stored by an Axona DacqUSB acquisition system. Spikes were amplified 3000-5000 times and bandpass-filtered between 0.8 and 6.7 kHz; local field potential (LFP) signals were amplified 1000 times and filtered between 0 and 475 Hz. Two light-emitting diodes (LEDs), one large and one small and separated by 5 cm, were attached to the headstage assembly to provide the position and the orientation of the rat’s head. The LEDs were imaged with an overhead camera at a sampling rate of 50 Hz.

Beginning 1 week after surgery, the animals were screened daily for single unit activity. Screening were made in a small box, outside the curtain. Tetrodes were lowered 50 p,m if no single unit activity was found, o r after a complete recording session. When a set of units was isolated, the animal was transferred to the experimental apparatus inside the curtained area and a sequence of five recording trials was run, as follows:

Trial 1 (Basline): The orientation of the box was random with respect to the outside room. Rats were placed in one of the compartments. For two RSC implanted rats, starting compartments were initially the same for every trial, but then switched to random. For the remaining two RSC implanted rats it was always random. For 10 minutes the animals freely moved between compartments foraging for cooked sweetened rice thrown in sporadically by the experimenter.

Trial 2 (Rotated baseline): As for Trial 1 but the box was randomly rotated by 45° or 90° or 180°, clockwise or counter-clockwise, to check local cue control of directional firing.

Trial 3 (Door closed odour 1): The apparatus was again arbitrarily rotated and the animal placed in one of the compartments (randomly chosen): the connecting door was closed so the rat was confined to that compartment for a five-minute trial. This was to see whether the odor cues alone were enough to reinstate the correct orientation of firing.

Trial 4 (Door closed odour 2): The apparatus was rotated and the rat recorded in the other compartment for five min.

Trial 5 (Baseline): A last standard 10-min door-open trial was run with the apparatus having the same orientation as in Trial 1.

Between trials, animals were removed from the apparatus and placed in a box in a random location outside the curtain for 2 minutes, in order to let the experimenter manipulate the box (rotation of the apparatus and/or closing or opening the door). The animal was then mildly disoriented, by rotating its holding box, before the next trial.

For some recording sessions, two ten-minute darkness trials were added after Trial 5. For the first of these (Trial 6 - Darkness), animals were not removed from the box after Trial 5, but the computer screen and ceiling lights were turned off before recording began. After this trial, the animal was removed, the box randomly rotated and the rat mildly disorientated before the last darkness trial (Trial 7 - Rotated Darkness).

### Data analysis

Spike sorting was performed manually using the graphical cluster-cutting software Tint (Axona) with the help of an automated clustering algorithm (Klustakwik 3.0; (Kadir et al., 2014)). Units having inter-spike intervals < 2 ms (refractory period), were removed due to poor isolation, as were cells with a peak firing rate <= 1 Hz. In order to prevent repeated recordings of the same cell over days, clusters that recurred on the same tetrodes in the same cluster space across recording sessions were only analysed on the first day.

The rat’s head direction was calculated for each tracker sample from the projection of the relative position of the two LEDs onto the horizontal plane. The directional tuning for each cell was obtained by dividing the number of spikes fired when the rat faced a particular direction (in bins of 6°) by the total amount of time the rat spent facing that direction. The peak firing rate was defined as the rate in the bin with the highest rate; the angle of this bin is the cell's preferred firing direction.

### Directionality analysis

The Rayleigh vector was used to assess directional specificity, with significant directionality being assigned to cells whose Rayleigh vector score exceeded the 99^th^ percentile of a control distribution. For the control procedure, for each cell’s Trial 1 data we shifted the time of every spike by a uniform amount, chosen randomly from between 20 seconds and the recording duration minus 20 seconds; this was repeated 400 times per cell (1090 cells × 400 = 436000 rotations). The 99^th^ percentile value came to 0.26, similar to that reported in previous HD cell studies (Bonnevie et al., 2013; Giocomo et al., 2014). Thus, all cells for which the Rayleigh vector length was greater or equal to 0.26 and firing rate exceeded 1Hz were classified as HD cells. The cells selected for the study were required to reach selection criteria in both trials 1 and 5.

Because of the unexpected observation of cells that fired in two opposing directions, we adapted the above directionality analysis by using an angle-doubling procedure in which the heading angle of each spike was multiplied by two (Batschelet, 1981): this has the effect of converting a bipolar distribution to a unimodal one, from which the population of directionally firing neurons could be selected using the 99^th^ percentile criterion computed as before and equal to 0.22.

We tested whether all 360° of compass heading were represented in the directional cells we recorded by referencing the firing direction of each cell to the lemon-compartment orientation in Trial 1. We then tested the non-uniformity of the distribution of angular distances with a Rayleigh test.

### Analysis of bipolar cells

#### Selection of bipolar cells

We selected bipolar HD cells by defining a “flip score” above which cells were considered bipolar, and below which they were unipolar. Flip score was calculated with an autocorrelation procedure, by rotating the polar firing rate plot in steps of 6° and calculating the correlation between the rotated and unrotated plots at each step. The bipolar pattern was apparent as a sinusoidal modulation of this autocorrelation with a peak at 180° (Supplementary Data Fig. 7). The flip score for each cell was defined as the difference between the correlations at the expected peak (180°) and the expected troughs (90°). For the entire population of directional cells this yielded two clusters (Fig. 2 and Supplementary Data Fig.7), the local minimum between which – value 0.6 – was used to separate bipolar from unipolar cells. Bipolar cells selected for the study were those which reached vector length and flip score criteria in trials 1 and 5.

#### Cell isolation analysis

Cell isolation analyses were performed to rule out the hypothesis that bipolar cells were really two co-recorded classic HD cells with opposing tuning curves.

1. Cluster space analysis (Supplementary Data Fig. 4a). This analysis sought to establish that the clusters from the spikes belonging to the two individual tuning curves of bipolar HD cells were no further apart than the clusters of clearly unipolar cells arbitrarily divided in two. This was based on scatterplots of the key waveform clustering parameter, which was peak-trough amplitude. For bipolar cells, we extracted two sub-clusters containing spikes emitted in the 180-degree range surrounding each tuning curve peak, found their center of mass (CoM) in the scatterplot cluster-space, and calculated the distance between the two CoMs. Results were compared to a control data set comprising the spikes from unipolar HD cells that had been randomly allocated to one or other of two sub-clusters; we then calculated the distance between the CoMs of these. This procedure was repeated 2000 times per cell. Data from the bipolar cells and the control data were compared with a two-sample *t*-test.
2. In a second analysis (Supplementary Data Fig. 4b), we calculated the probability that failure to separate the clusters of unipolar HD cells would have resulted in tuning-curve pairs that just happened to be 180 degrees apart. To do this we randomly (with replacement) selected pairs of HD cells from the total pool, 10000 times, and for each pair we cross-correlated the tuning curves and derived the angle of the highest correlation. Then, we calculated the percentage of cells with an angular distance at 180±12° (corresponding to two bins of 6°) and compared this, using a Chi-squared test, with the observed data.
3. In the last analysis, we computed the Pearson’s correlation coefficient between firing rates for the two tuning curves (Supplementary Data Fig. 4c) – if these were from different cells then the rates should be no more correlated than those of any two randomly selected cells. The correlation value was compared to control correlations generated by randomly selecting, with replacement, 10 000 pairs of HD cells.

#### Directionally color-coded spatial spike maps

Directional color-coded spike maps were constructed in order to visually display the spatial distribution of directional activity. We first defined a colour map with a circular gradient of colour ranging in 90-deg bins from blue to green to red to yellow (Fig. 1c). For each cell, we aligned the direction of blue in the colour map with the peak firing bin in Trial 1, with all other directions being colour accordingly, and assigned each spike the appropriate colour based on its associated head direction. We then generated a spatial spike plot by smoothing the path of the rat in 20 ms bins and overlying it at the appropriate places with the color-coded spikes.

#### Within-compartment bipolar cell analysis

Some neurons had bipolar tuning curves even in individual sub-compartments of the apparatus and so we investigated whether these behaved like unipolar cells that periodically reversed their tuning curve direction, or more like cells with simultaneous, opposing tuning curves. Analysis was performed on the two tuning curves for the whole apparatus, and also for each individual compartment on the baseline trial. We identified epochs in the recording trial when the rat faced continuously in the direction (+/− 45 deg) of one or other tuning curve, and then within these epochs we computed the firing rate (no. spikes divided by epoch duration), and firing rate spread (standard deviation of firing rates across epochs).

#### Directional activity in darkness and in the open field.

Directionally modulated cells were recorded in darkness so as to determine whether activity was maintained between light and dark trials. For each cell we computed (1) the vector length and the flip score as described previously, and (2) the Pearson correlation coefficient of each cell’s directional firing rates between continuous light-dark trials 5 and 6, and between the two dark trials 6 and 7, and compared them using a Kruskal-Wallis non-parametric test.

Nine bipolar cells were additionally recorded in a 120 cm × 120 cm open-field after 5 experimental recording sessions. For each bipolar cell, we calculated the vector length and the flip score to investigate whether they share directional properties found in the two compartment apparatus.

#### Spatial rate maps and spatial correlation analysis

Spatial rate maps were generated by dividing the two-compartment apparatus into an array of 40 × 40 square bins, each 3 × 3 cm in size. Spikes per bin were divided by time spent in that bin to provide a firing rate (Hz). Smoothed firing rate maps were then generated following Spiers et al. (Spiers et al., 2015) by using a boxcar procedure in which the firing rate in each bin was replaced by that of the mean of itself plus the immediately surrounding bins (8 for central bins, 5 for edge bins and 3 for corner bins). In both raw and smoothed rate maps, pixels that were not visited by the rat were displayed in white. The firing rate of the cell is color-coded from low (light blue) to high (dark red).

To estimate the stability of the spatial activity, pixel-by-pixel cross-correlations were made on the unsmoothed data from the two compartments – the first using the original maps, and the second with one compartment rotated by 180°, in order to see whether spatial patterns reversed in the same way that the directional patterns of bipolar cells did.

### LFP/spiking characteristics and movement correlates

For recordings from the three brain areas we looked at local field potential (LFP) theta frequency and power, while for the six cell types (PoS HD, ADN HD, RSC unipolar, RSC between-compartment bipolar and RSC between-compartment unipolar) we looked at spiking characteristics including peak firing rate, inter-spike interval (ISI) histogram. For both LFP and spiking we looked at theta modulation, and correlation of firing rate with running speed and angular head velocity.

For the ISI analysis, a histogram of ISIs with 2ms bins was generated for each cell type and the peak was taken as the centre of the bin with the highest count. Inspection of the ISI histograms suggested different decay times for the different cell types so we calculated decay time by fitting, to the histogram, a one-term exponential decay function of the form y = a*exp(b*x) from the peak to peak + 1 second, using the *fit* function from MATLAB's Curve Fitting Toolbox. Time to half-peak was then taken as the time taken for the exponential fit to decay to half the peak value.

For retrosplenial bipolar within-compartment, bipolar between-compartment, and unipolar cells, ISIs were also considered only for periods of time where HD was within +/− 45 degrees of the preferred firing direction of the cell. For unipolar HD cells, this was taken as the location of peak firing rate in the polar plot. For bipolar HD cells, this same method was used to find one preferred firing direction and then the other peak was found using a circular autocorrelation. For the exponential curve-fitting, the lowest-spiking 25% of all cells were excluded (those with fewer than 145 spikes occurring within +/− 45 degrees of the peak direction for that cell). One-way analysis of variance did not reveal significant differences between cell types when considering exponential fitting parameters a [F(2,157) = 0.1; NS] and b [F(2,157) = 1.57; NS], nor when considering rise time to peak [F(2,157) = 0.25; NS] and decay time to half-peak [F(2,157) = 1.33; NS].

#### LFP theta and speed-theta correlation

Running speed was related to LFP theta frequency in the manner of Jeewajee et al. (Jeewajee et al., 2008). The LFP signal was filtered with the 251-tap Blackman windowed band-pass sine cardinal filter in the theta range (7-11Hz). The analytic signal was determined used the Hilbert transform, with the instantaneous theta frequency then taken as the difference in phase between each time point (averaged over 5 consecutive values). This compensated for sampling of position at 50Hz and LFP signal at 250Hz, allowing comparison at 50Hz.

The relationship between speed and frequency was taken as the *z* correlation coefficient derived from the *r* correlation via the Fisher transformation. A regression line was then fitted between theta frequency and running speed in the range 4-30cm/s. The slope of this regression line was then reported.

The power spectrum was created by zero-padding the LFP sequence to the next highest power of 2 and taking the fast Fourier transform (FFT). The square-modulus of each Fourier frequency coefficient gave the signal power at that frequency. The power spectrum was smoothed with a Gaussian kernel (width 2Hz, sd 0.1875Hz). Theta frequency was taken as the frequency at which the power spectrum peaked within the theta range. Theta power was taken as the mean power in the theta band (7-11Hz) divided by the mean power in the delta band (1-4Hz).

#### Intrinsic frequency

Intrinsic frequency for a given cell was estimated by taking the power spectrum of its spike train autocorrelogram. The autocorrelogram was first truncated to the first 0.5 s and zero padded to 2^16^ elements. Its power spectrum was then found, and the theta frequency was taken as the peak of this power spectrum in the range 7-11 Hz. Theta power was taken as the mean power in the theta band (7-11Hz) divided by the mean power in the delta band (1-4Hz).

#### Linear and angular speed

Both linear and angular speeds were related to firing rate. Linear speed, in cm/s, for each camera frame was calculated as the difference in position of the rat across successive video frames divided by the time between frames (20ms with a 50Hz camera sample rate). Angular speed, in deg/s, was calculated as the change in head direction rat across successive video frames divided by the time between frames.

Linear speeds in the range 2 to 30cm/s were binned in 2cm/s intervals, whilst angular speeds in the range −500 deg/s to 500deg/s were binned in 2 deg/s intervals. For each bin, the number of spikes and dwell time were used to calculate a firing rate. Any bin with a dwell time of < 0.5s was excluded from later analysis. A regression line between bin means and firing rate for each bin was fitted using the MATLAB curve-fitting toolbox.

The relationship between linear or angular speed and firing rate was only considered when the animal’s head direction was within 45 degrees either side of the preferred firing direction of the cell. For unipolar HD cells, this was taken as the location of peak firing rate in the polar plot. For bipolar HD cells, this same method was used to find one preferred firing direction. A circular autocorrelation was used to find the location of maximum correlation of the bipolar cell polar plot with itself. The second preferred firing direction was then taken as being separated from the first by the distance between circular autocorrelogram peaks.

### Histology

At the completion of the experiment, rats received an overdose of pentobarbital and were perfused intracardially with 0.9% saline followed by 4% formaldehyde. The brains were removed, stored 1 day in formaldehyde, followed by 30% sucrose solution, and finally frozen with dry ice. 30- μm-thick coronal sections for ADN and RSC animals and sagittal sections for PoS animals were mounted on glass slides and stained with cresyl violet or thionine. The position of the tips of the electrodes were determined from digital pictures, acquired with an Olympus microscope (Olympus Keymed, Southend-on-Sea, U.K.) and imported into an image manipulation program (Gimp 2.8, distributed under General Public License).

Recording sites were determined by measuring backwards from the deepest point of the track, and identifying with the help of the rat brain atlas from Paxinos and Watson (Paxinos and Watson, 2007).

